# Maternal factor PABPN1L is essential for maternal mRNA degradation during maternal-to-zygotic transition

**DOI:** 10.1101/2020.08.20.258830

**Authors:** Ying Wang, Tianhao Feng, Xiaodan Shi, Siyu Liu, Zerui Wang, Xin Zhang, Jintao Zhang, Shuqin Zhao, Junqiang Zhang, Xiufeng Ling, Mingxi Liu

**Author notes:** These authors contributed equally to this work (Ying Wang, Tianhao Feng, Xiaodan Shi). Corresponding authors: Mingxi Liu, E-mail address, Xiufeng Ling.

## Abstract

Infertility affects 10% - 15% of families worldwide. However, the pathogenesis of female infertility caused by abnormal early embryonic development is not clear. We constructed a mouse model (Pabpn1l -/-) simulating the splicing abnormality of human PABPN1L and found that the female was sterile and the male was fertile. The Pabpn1l -/- oocytes can be produced, ovulated and fertilized normally, but cannot develop beyond the 2-cell stage. Using RNA-Seq, we found a large-scale upregulation of RNA in Pabpn1l -/- MII oocytes. Of the 2401 transcripts upregulated in Pabpn1l-/- MII oocytes, 1523 transcripts (63.4%) were also upregulated in Btg4 -/- MII oocytes, while only 53 transcripts (2.2%) were upregulated in Ythdf2 -/- MII oocytes. We documented that transcripts in zygotes derived from Pabpn1l -/- oocytes have a longer poly(A) tail than the control group, a phenomenon similar to that in Btg4-/- mice. Surprisingly, the poly(A) tail of these mRNAs was significantly shorter in the Pabpn1l -/- MII oocytes than in the Pabpn1l +/+. These results suggest that PABPN1L is involved in BTG4-mediated maternal mRNA degradation, and may antagonize poly(A) tail shortening in oocytes independently of its involvement in maternal mRNA degradation. Thus, PABPN1L variants could be a genetic marker of female infertility.

## Introduction

Infertility affects the happiness of millions of families. It is reported that 10-15% of couples of childbearing age are facing infertility ^1, 2^. The emergence and vigorous development of assisted reproductive technology have benefited many infertile patients. However, in vitro fertilization-embryo transfer (IVF-ET) is frequently accompanied by retardation in embryo development. Addressing this problem and improving the quality of preimplantation embryos is an urgent challenge for assisted reproductive technology.

In mammals, the time of the arrest of embryo development varies with species. Early human embryos cultured in vitro are prone to the block of development at the 4- or 8-cell stage^3^, while in mouse embryos, it takes place typically at the 2-cell phase. This time point coincides with zygotic gene activation (ZGA), a critical event in early embryonic development, suggesting that ZGA may be closely related to embryonic development arrest ^4–6^. Since transcription is silent in oocytes before ZGA^7, 8^, embryonic development is primarily regulated by maternal factors, including mRNA and protein, accumulated and stored in the cytoplasm of mature oocytes during oogenesis ^9, 10^. After ZGA, these maternal factors activate the embryonic genome and initiate transcription to express new genes and proteins, and a cascade of reactions occurs to maintain the development of the embryo ^11^. Maternal mRNA begins to be degraded after oocyte maturation and is replaced by new mRNA synthesized after zygote genome activation. This process is accompanied by the clearance of maternal proteins and is called maternal to zygotic transition (MZT)^10^. In the case of abnormal clearance of maternal mRNAs, the development of early embryos is blocked^12^. Therefore, identification of the mechanisms governing the clearance of maternal mRNAs clearance may provide a novel theoretical basis for understanding the causes of preimplantation embryo development retardation and improve the success rate of IVF-ET.

Due to the inhibition of the transcriptional activity before the genome of the zygote is activated, the expression of protein mainly depends on mRNA stored in the egg ^8, 13^. These mRNA molecules are stable, with a half-life of 8-12 days ^14, 15^. They participate in the processes of meiosis and cleavage after fertilization. Generally, the mRNA synthesized in the nucleus contains poly(A) tail, which is about 200-250 bp in length^16^. Poly(A) tail is closely related to mRNA stability, transport, and protein translation ^17^. The poly(A) tails are not genetically encoded but are formed by the cleavage of specific sites of precursor mRNA (pre-mRNA) and polyadenylation ^18, 19^. It has been demonstrated that mRNA stored in oocytes and early embryos is regulated by several RNA binding protein complexes ^20, 21^. For example, the RNA binding protein CPEB1 (cytoplasmic polyadenylation element-binding protein 1) interacts with other regulatory proteins, such as SYMPK, CPSF, and GLD2, to regulate the formation of mRNA poly(A) tail, which is essential for the maturation of oocytes ^21–23^. During MZT, the deadenylation of mRNA poly(A) tail is mediated by the BTG4-EIF4E-EIF4G-mRNA complex. The absence of BTG4 leads to embryo arrest ^12^. Similar phenotypes were also found in mice with the knockout of an m6A reader, Ythdf2. The oocytes of Ythdf2-/- mice exhibited a 2-cell block after fertilization, which resulted in the upregulation of mRNA in MII oocytes^24^. These studies suggest that RNA-binding proteins may play an important role in early embryonic development.

Previous studies have documented that poly(A)-binding proteins (PABPs) function as cis-acting effectors of specific steps in the polyadenylation, mRNA export, translation, and turnover of the transcripts to which they are bound ^17^. PABPs can be divided into the cytoplasmic type and nuclear type. Cytoplasmic PABPs are highly conserved and contain four RNA recognition motifs (RRM) differing in their affinity to RNA poly(A) tails. Among cytoplasmic PABPs, ePAB (embryonic poly(A)-binding protein) is necessary for the maturation of mouse and Xenopus oocytes^25^. Nuclear PABPs, such as PABPN1 and PABPN1L, contain one RRM ^17^. PABPN1 can bind 11 to 14 adenylate residues in mammalian cells ^26^ and interacts with the cleavage and polyadenylation specificity factor (CPSF) to activate the synthesis of RNA poly(A) tail ^27^. Also, PABPN1 is involved in mRNA splicing ^28^ and may be related to a viral protein capable of blocking the nuclear transport of mRNA ^29^. The homolog of PABPN1L, ePABP2, which belongs to nuclear PABPs, is expressed in oocytes and early embryos of Xenopus ^30^. PABPN1L can bind the AAAA nucleic acid sequences in human cell lines, but it is not clear what role it plays in organisms ^31^.

In the present study, a mouse model (Pabpn1l -/-) was constructed to mimic the splicing abnormality of PABPN1L in humans. We found that in this mouse model, the female was sterile while the male was fertile. Pabpn1l -/- mice can produce oocytes, ovulate, and fertilize normally, but cannot develop beyond the 2-cell stage. Through RNA-seq, we documented that RNA upregulation began to appear in MII oocytes of Pabpn1l -/- mice and remained significant also after fertilization. In comparison with the Btg4 knockout mice and Ythdf2 knockout mice, 63.4% of genes upregulated in the MII phase were also upregulated in Btg4 -/- MII phase oocytes, and only a few of the upregulated genes upregulated in the Ythdf2 -/- oocytes. The poly(A) tail (PAT) assay demonstrated that the poly(A) tail of the upregulated transcripts in Pabpn1l -/- zygote was longer than in Pabpn1l +/+. These results suggest that PABPN1L is involved in the degradation of maternal mRNA, and its activity overlaps with BTG4. Interestingly, we found that the poly(A) tail of mRNA in MII oocytes of Pabpn1l -/- mice was significantly shortened, and even shorter than in Pabpn1l +/+. This phenomenon is different from that present in Btg4-/- MII oocytes. These results suggest that PABPN1L, as a maternal factor, plays an essential role in the degradation of maternal mRNA, and may inhibit deadenylation in MII oocytes, which is independent of maternal mRNA degradation.

## Results

### Pabpn1l is a conserved maternal expression gene

The sequence of PABPN1L protein was highly conserved in vertebrates (Figure 1A). qRT-PCR documented that Pabpn1l mRNA was present in oocytes and early embryos. The expression was highest in the germinal vesicle (GV) phase and decreased gradually with oocyte maturation and fertilization (Figure 1B). This trend is consistent with the characteristics of maternal factors, which are highly expressed before the zygote genome is activated. The expression of Pabpn1l mRNA in the ovarian tissue was significantly higher in other tissues (Figure 1B), which may be due to the presence of a small number of GV oocytes in the ovary. Since the transcription of Pabpn1l is the highest in the GV phase, we injected PABPN1L-EGFP mRNA into the cytoplasm of GV phase oocytes to detect the localization of PABPN1L-EGFP in oocytes. In contrast to the cytoplasmic localization of ePABP2 in GV phase oocytes of Xenopus laevis, PABPN1L-EGFP was enriched in the nucleus of oocytes and relocated to the oocyte cytoplasm after nuclear membrane rupture (Figure1C).

**Figure 1:**
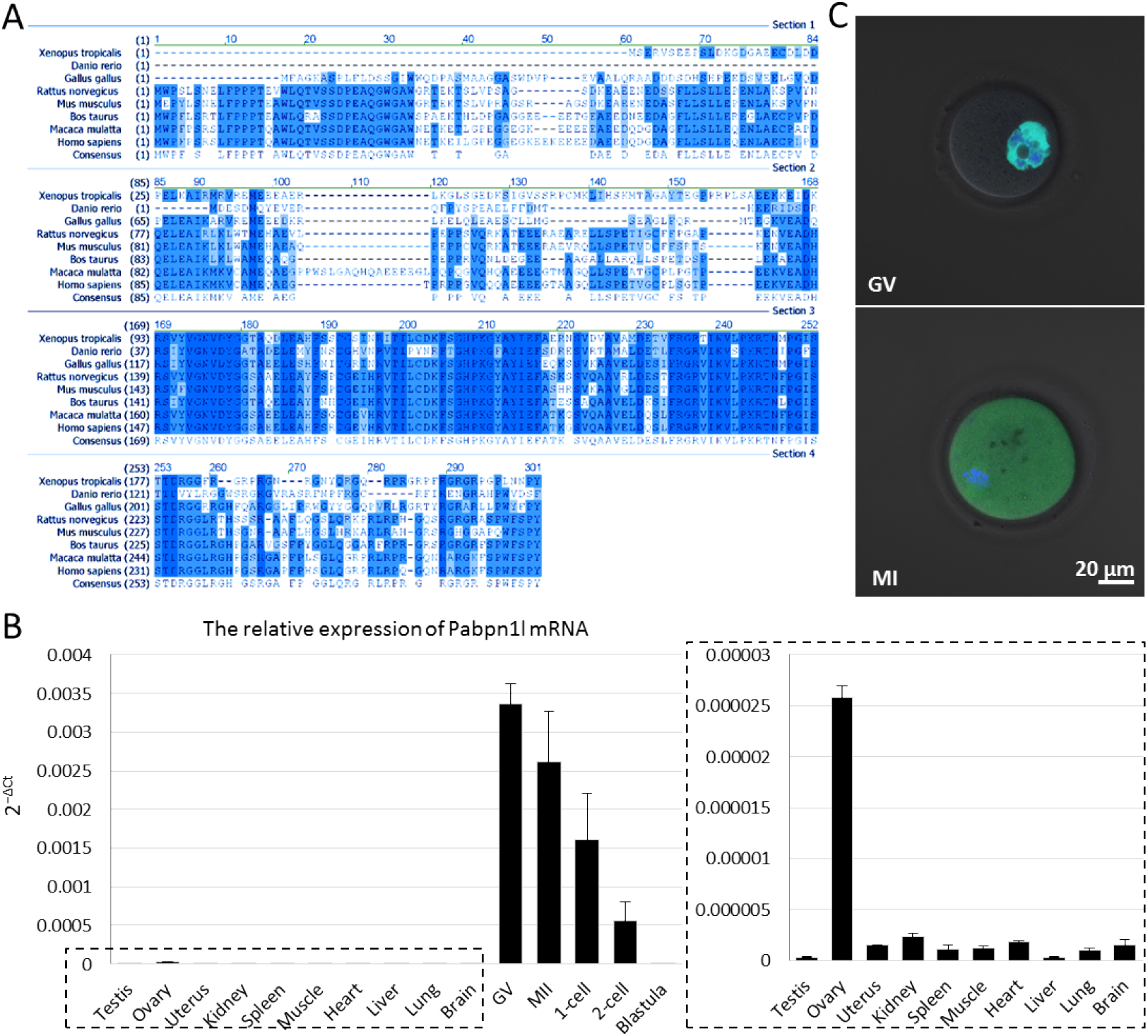
PABPN1L is a conserved maternal protein. (A) Sequence alignment of PABPN1L proteins from several vertebrates. Conserved amino acid sequences are highlighted in blue. (B) Quantitative RT–PCR results showing relative expression levels of Pabpn1l in mouse tissues (indicated in the dotted box), oocytes, and preimplantation embryos (n= 3). (C) PABPN1L-EGFP located in the nucleus of GV oocytes and relocated to the cytoplasm in MI phase oocytes after injection of PABPN1L-EGFP mRNA into the cytoplasm of GV phase oocytes.

### Abnormal splicing of Pabpn1l mRNA leads to female infertility in mice

Through the analysis of the gnomAD data (https://gnomad.broadinstitute.org), we found that there were nine single-nucleotide variants (SNVs) on intron1 that might cause splicing abnormality. Among them, rs759387263, rs537683283, and rs7524277449 close to Exon2 were recurrently detected, with rs759387263 having as many as 26 carriers (Figure 2 and Table 1). These SNVs may affect the normal function of PABPN1L in humans. Using CRISPR/cas9, we constructed the mutation model Pabpn1l -/- (Figure 3A) at the junction of intron1 and exon 2 of Pabpn1l. In the ovary of Pabpn1l -/- mice, intron1 retention of Pabpn1l mRNA was observed (Figure 3B), and most of the abnormally spliced mRNA was eliminated in vivo (Figure 3C). The fertility test demonstrated Pabpn1l-/- female infertility, but normal male fertility (Figure 3D).

**Figure 2:**
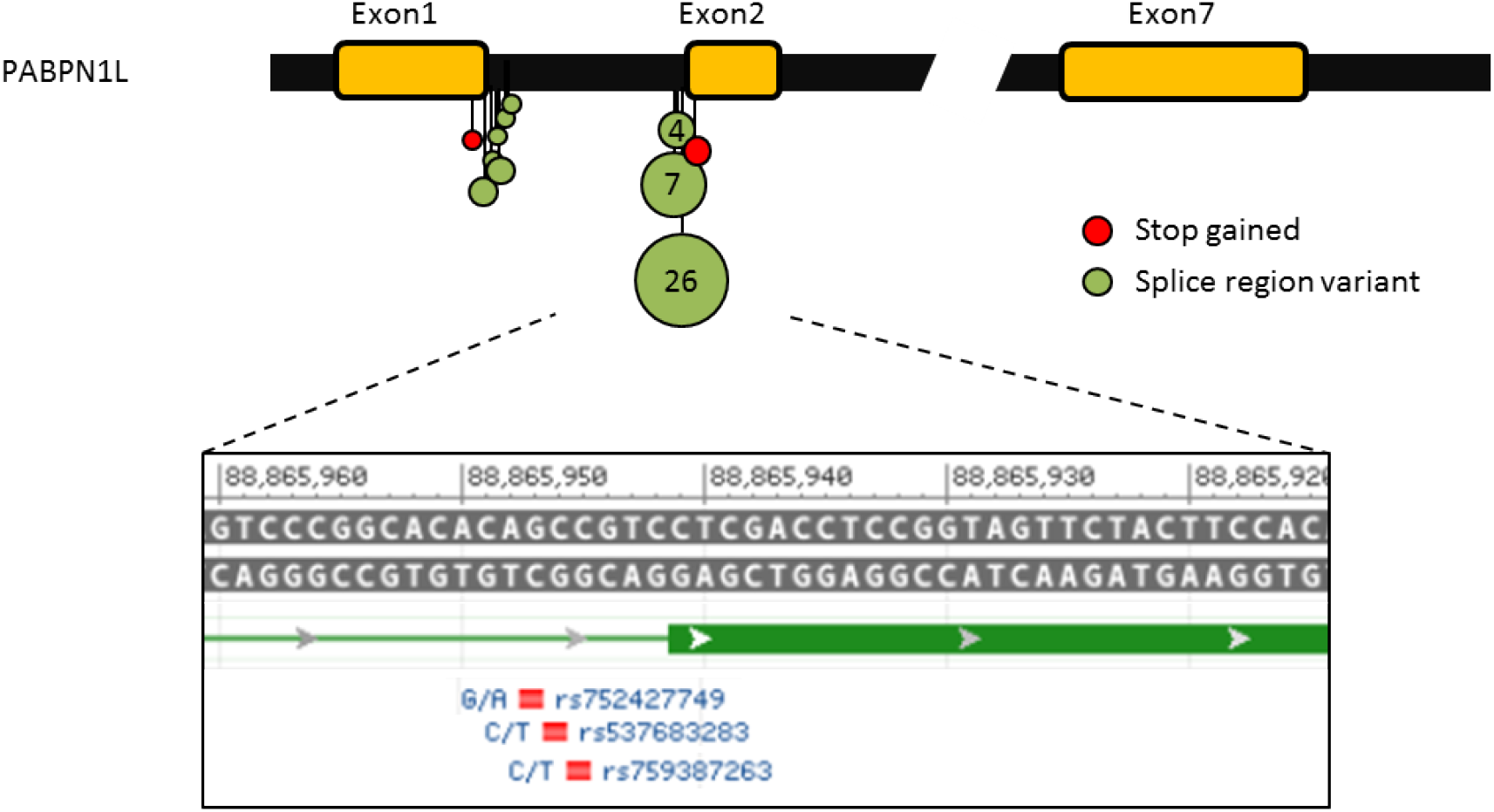
Schematic of SNVs in exon1-intron1 junction and intron1-exon2 junction of PABPN1L. The size of the circle indicates the number of recurrences of SNVs. Rs759387263, rs537683283, and rs7524277449 close to Exon2 were recurrently detected.

**Table.1.**
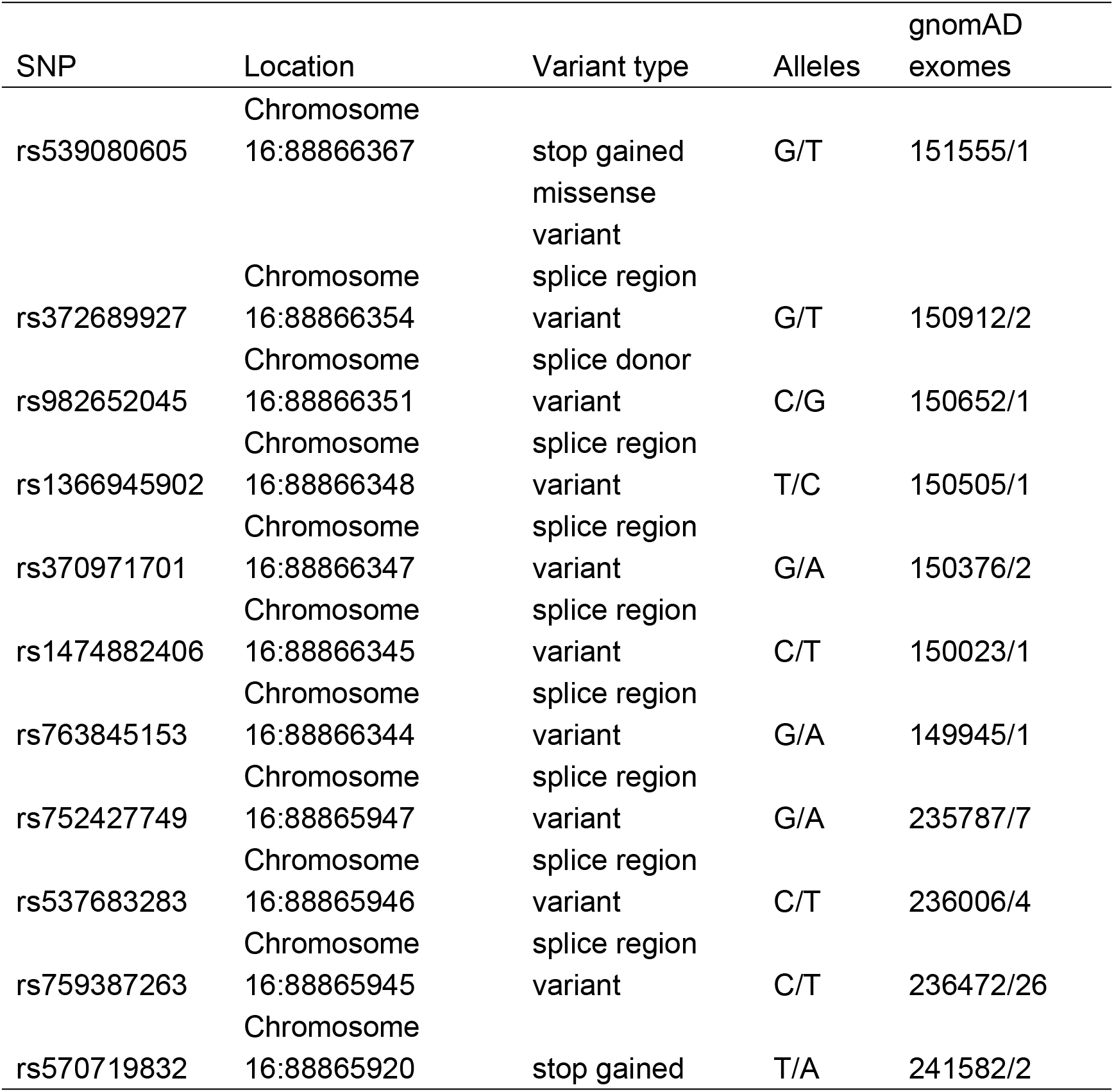
Potential severe variation at the junction of exon 1 and exon 2 of PABPN1L

**Figure 3:**
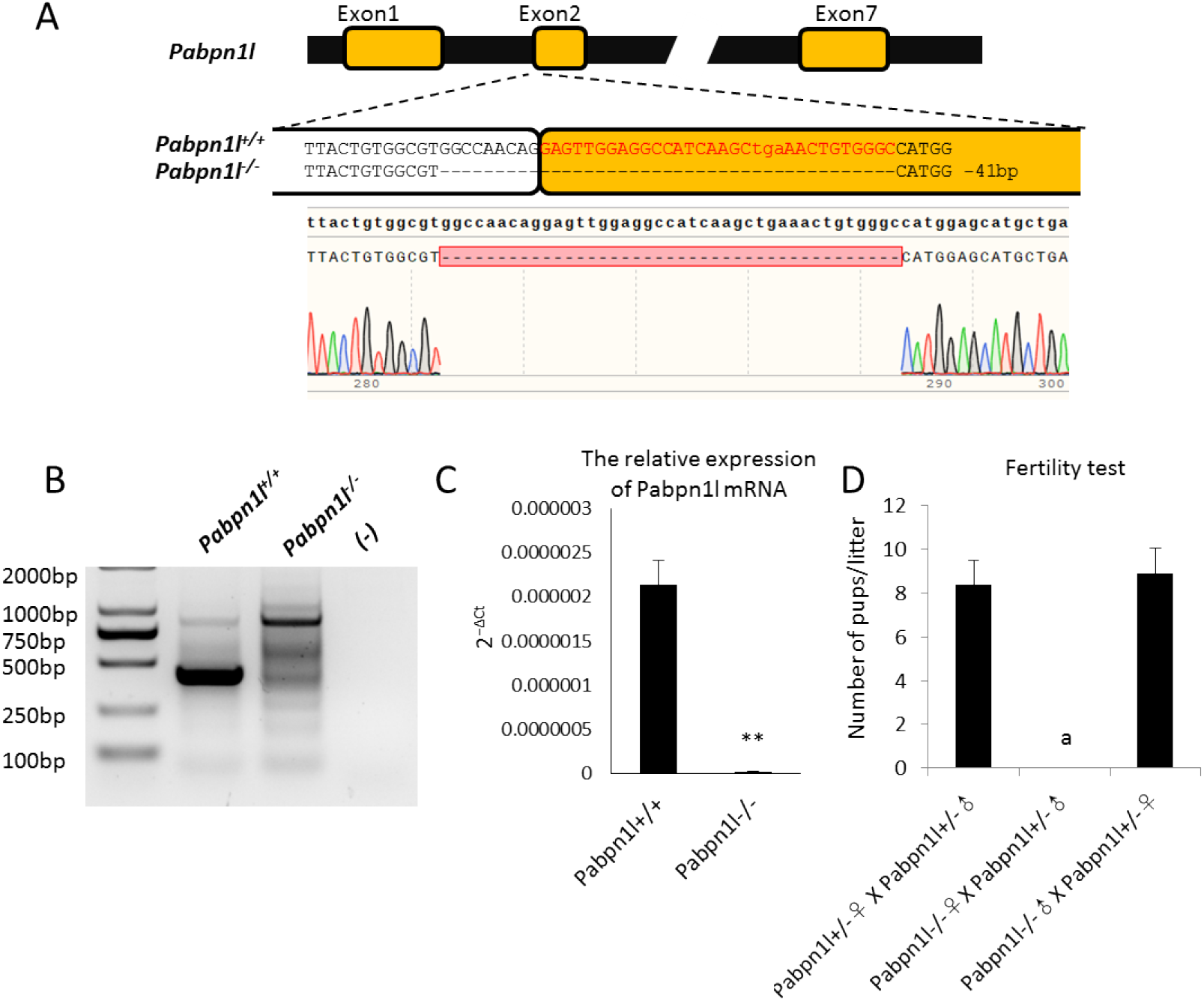
Abnormal splicing of Pabpn1l mRNA in Pabpn1l -/- mice leads to female infertility. (A) A 41-bp deletion of intron1-exon2 junction of Pabpn1l was detected in Pabpn1l -/- mice by Sanger sequencing. (B) RT-PCR products using primers targeted to exons 1– 3 of the Pabpn1l transcripts of the ovary. The 41-bp deletion of intron1-exon2 junction leads to intron1 retention in Pabpn1l mRNA. (C) Quantitative RT–PCR results showing relative expression levels of Pabpn1l in the ovary of Pabpn1l +/+ and Pabpn1l -/- mice. Most of the abnormally spliced mRNA was eliminated in Pabpn1l -/- ovary. (D) Pabpn1l -/- females are fertile. Bar graph showing the average number of pups/litter (n = 8).

### PABPN1L is a maternal factor essential for maternal-to-zygotic transition

To further evaluate the specific link between Pabpn1l deletion and female reproductive ability, we analyzed ovarian sections of adult Pabpn1l -/- mice. The ovarian morphology was generally normal, and all levels of follicles were visible (Figure S1A). There was no difference in the number of oocytes between Pabpn1l -/- and Pabpn1l +/+ mice (Figure S1B). The ratio of GV stage oocytes cultured to germinal vesicle breakdown (GVBD) was normal (Figure S1C), as was the emission rate of the first polar body (Figure S1C). Using in vitro fertilization (IVF), we found that Pabpn1l-/- oocytes could be fertilized normally, and female and male pronuclei were formed (Figure S2). However, the development of fertilized eggs from the 1-cell to 2-cell stage was blocked (Figure 4A and B).

**Figure 4:**
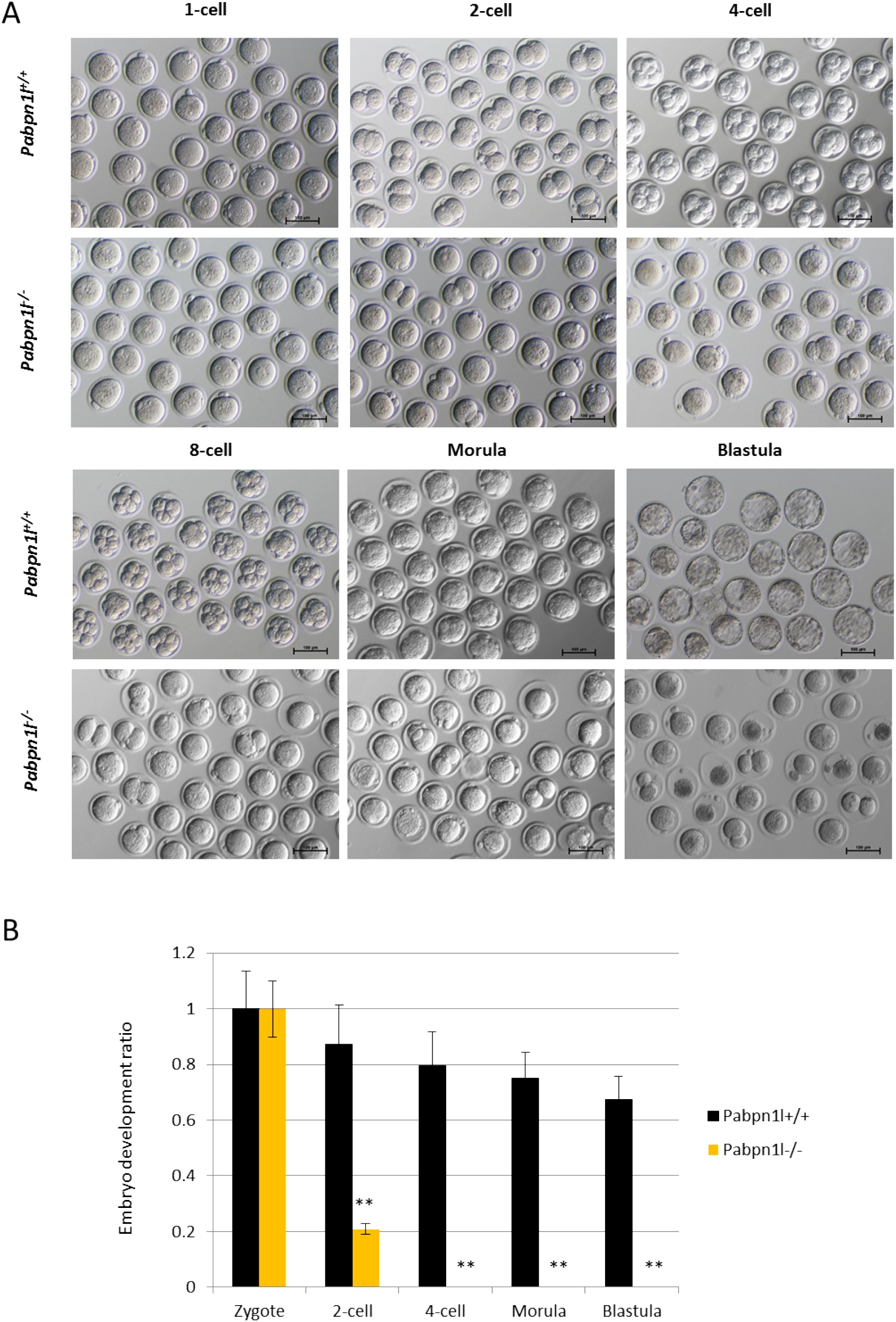
PABPN1L is essential for maternal-to-zygotic transition. (A) Representative images of the embryos obtained by in vitro fertilization and culture of Pabpn1l -/- oocytes and Pabpn1l +/+ oocytes at the indicated stages. Scale bar = 100 μm. (B) Developmental ratio of embryos obtained in vitro fertilization and culture of Pabpn1l -/- oocytes and Pabpn1l +/+ oocytes at the indicated stages.

In addition to the Pabpn1l-/- model, we generated another mouse model based on gene trapping strategy, Pabpn1l tm1a/tm1a (Figure 5A), and found that Pabpn1l mRNA was absent in the ovary of these mice (Figure S3). Female Pabpn1l tm1a/tm1a mice were infertile (Figure 5B). The ovarian morphology of Pabpn1l tm1a/tm1a was normal, and the number of ovulated oocytes was similar to that of wild-type mice (Figure S4). In Pabpn1l tm1a/tm1a mice, the oocytes at the MII stage could be normally fertilized, but the development of fertilized eggs from the 1-cell stage to 2-cell stage was blocked (Figure 5C), resembling the Pabpn1l -/- phenotype. Based on the phenotype of Pabpn1l -/- and the expression pattern of Pabpn1l, we conclude that Pabpn1l is a maternal factor.

**Figure 5:**
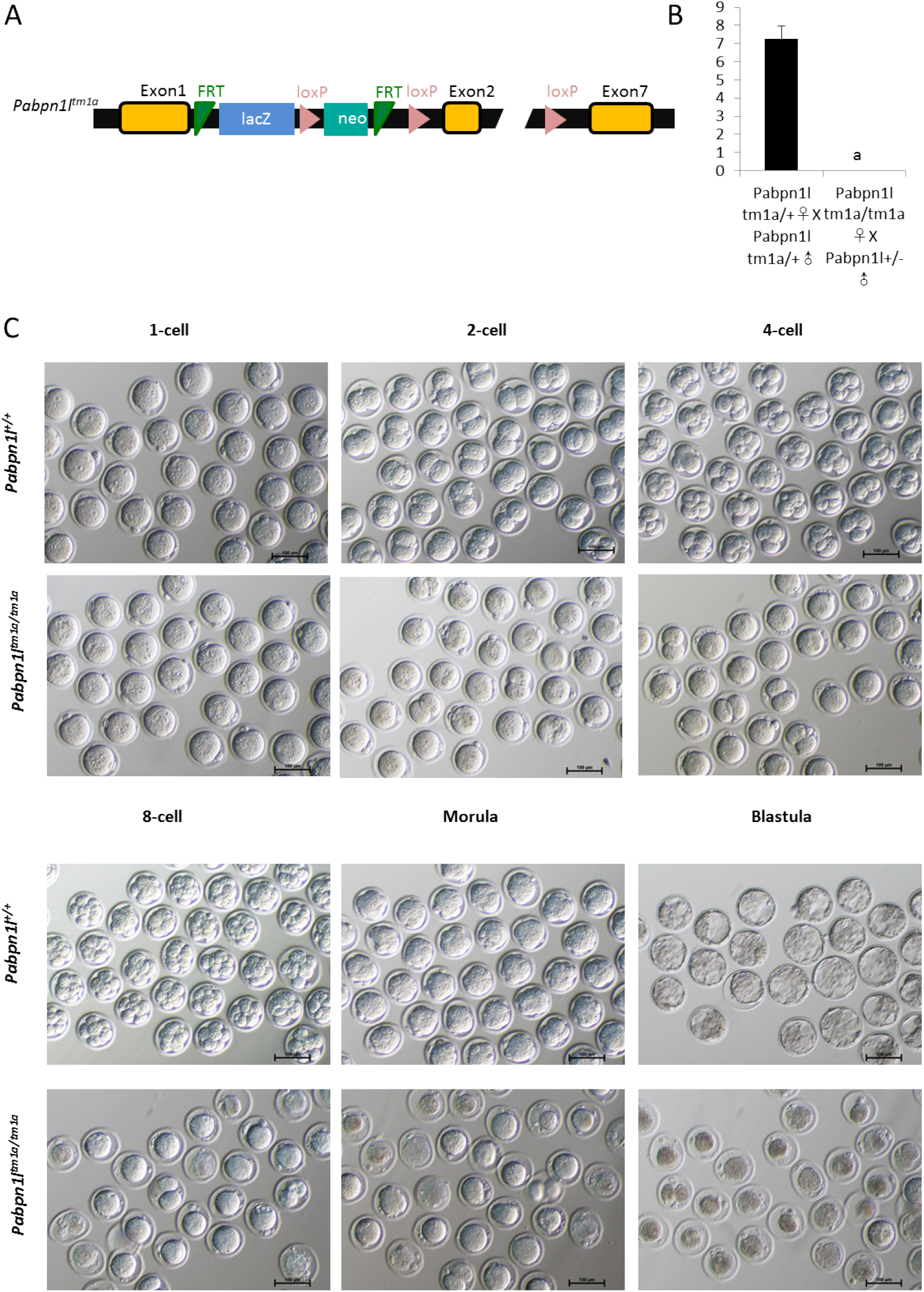
Female Pabpn1l tm1a/tm1a mice were infertile. (A) Schematic of the Pabpn1l tm1a allele. (B) Pabpn1l tm1a/tm1a females are fertile. Bar graph showing the average number of pups/litter (n = 7). (C) Representative images of the embryos obtained by in vitro fertilization and culture of Pabpn1l -/- oocytes and Pabpn1l +/+ oocytes at the indicated stages. Scale bar = 100 μm.

### PABPN1L is essential for maternal mRNA degradation during maternal-to-zygotic transition

Since PABPN1L is capable of binding the poly(A) tail, we raised the possibility that it participates in the process of post-transcriptional regulation of maternal mRNA. We investigated the mRNA changes in GV oocytes, MII oocytes, and fertilized eggs using RNA-seq (Figure 6A-C). There were 2401 upregulated and 398 downregulated transcripts in Pabpn1l-/- MII oocytes (Figure 6B). This large-scale upregulation of mRNAs persisted after IVF of Pabpn1l-/- MII oocytes with Pabpn1l +/+ sperm (Figure 6C). Subsequently, we compared the transcriptome data of Pabpn1l -/- MII oocytes with the published Btg4 -/- and Ythdf2 -/- data^12, 24^. Of the 2401 transcripts upregulated in Pabpn1l -/- MII oocytes, 1523 transcripts (63.4%) were also upregulated in Btg4 -/- MII oocytes, while only 53 transcripts (2.2%) were upregulated in Ythdf2 -/- MII oocytes (Figure 6D). Also, 1265 transcripts (62.7%) were upregulated in the transcriptome of Btg4-/- MII oocytes. Gene Ontology (GO) analysis revealed that the transcripts upregulated in Pabpn1l -/- and Btg4 -/- MII oocytes were enriched in mitochondrial and ribosomal pathways, suggesting that energy metabolism and protein translation need to be inhibited during MZT (Table S1). Btg4 is involved in the process of the degradation of maternal mRNA by promoting its deadenylation ^12^. The above results suggest that PABPN1L participates in BTG4-mediated maternal mRNA degradation.

**Figure 6:**
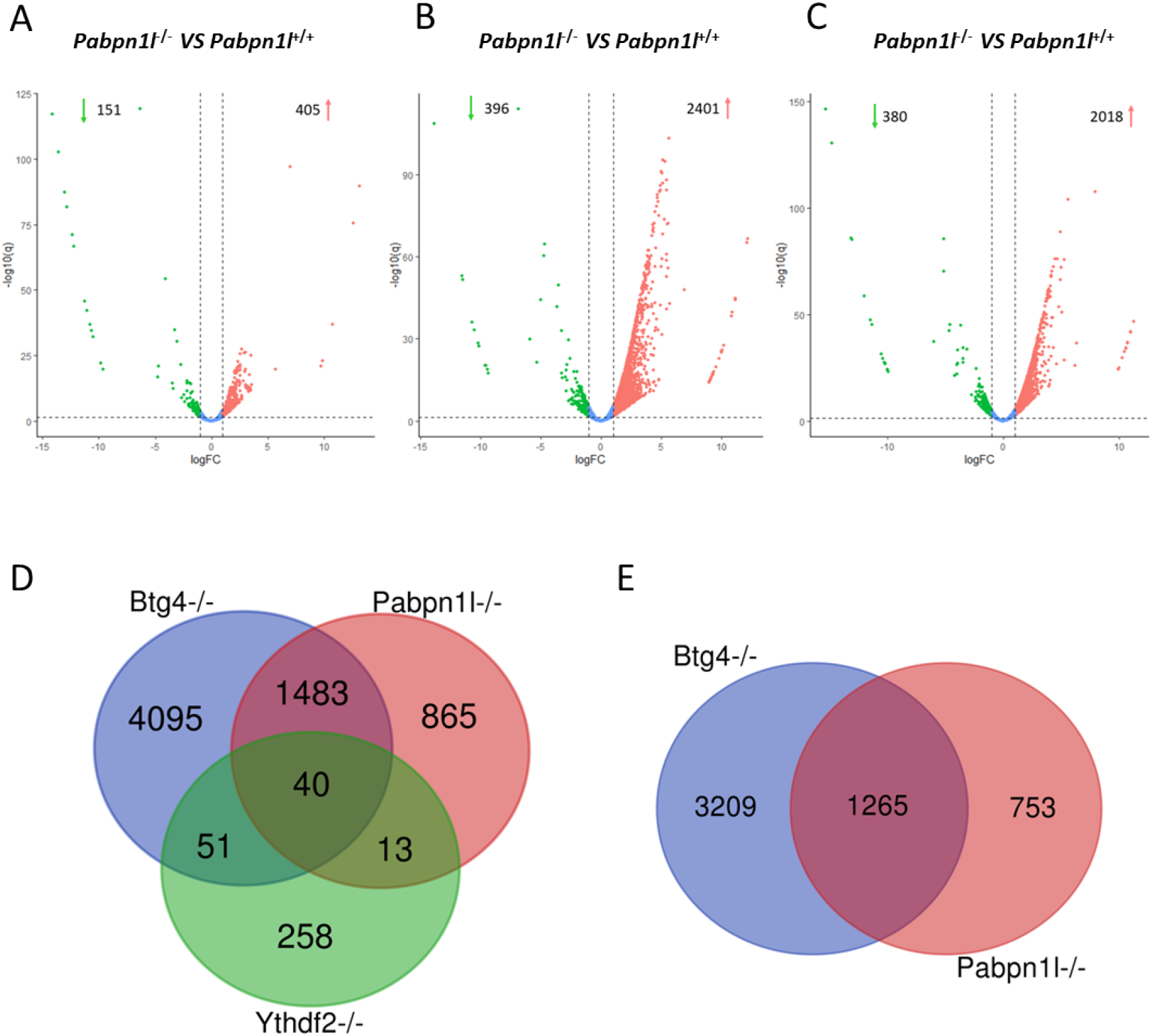
PABPN1L may be involved in the BTG4-mediated degradation of maternal mRNA. Volcano plots comparing the transcripts of GV oocytes (A), MII oocytes (B), and zygotes (C) from Pabpn1l -/- and Pabpn1l +/+ females. Transcripts that increased or decreased by more than 2-fold in Pabpn1l -/- deleted GV oocytes, MII oocytes or zygotes are highlighted in red and green, respectively. (D) Venn diagrams showing the overlap of transcripts that are significantly increased in MII oocytes from Pabpn1l-/-, Btg4-/-, and Ythdf2-/- mice. (E) Venn diagrams showing the overlap of transcripts that are significantly increased in zygotes from Pabpn1l-/- and Btg4-/- mice.

### Deletion of Pabpn1l leads to deadenylation of mRNA in MII stage oocytes, which is independent of maternal mRNA degradation

To investigate whether Pabpn1l mediates deadenylation of maternal mRNA, we selected 4 transcripts from the up-regulated transcripts in Pabpn1l -/- MII oocytes using the PAT assay. These four transcripts, Timm9, Eloc, Elobl, and Fbxw13, remained upregulated trend after fertilization, consistent with the trend of Btg4 -/- MII oocytes and zygotes. These four transcripts had longer poly(A) tail in zygotes derived from Pabpn1l -/- oocytes than the control group (Figure 7A and B), which is similar to the phenomenon in Btg4 -/- mice^12^. Interestingly, we found that the poly(A) tail of these mRNA was significantly shorter in the Pabpn1l -/- MII oocytes than in the control group. These transcripts did not degrade after the shortening of poly(A) tail, but accumulated in MII oocytes. Then, we selected a transcript Cdc123 that was not changed in Pabpn1l -/- MII oocytes. We found that the poly(A) tail of Cdc123 was also shortened in Pabpn1l -/- MII oocytes, and similar phenomena were observed in Gapdh transcripts used as an internal reference. Therefore, we hypothesize that PABPN1L inhibits the shortening of poly(A) tail in MII oocytes, but this function is independent of BTG4-mediated mRNA degradation.

**Figure 7:**
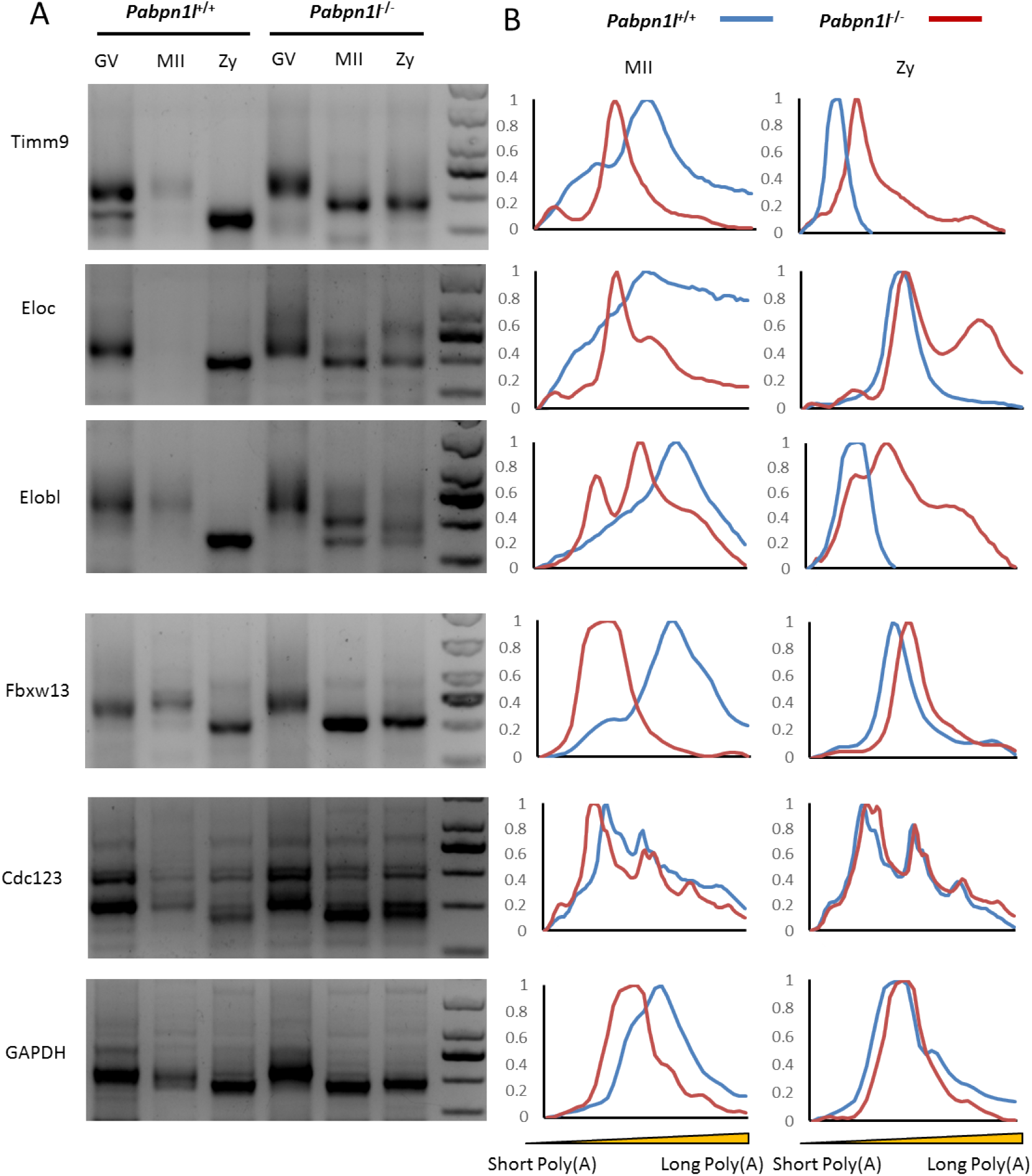
Poly(A) tail length of transcripts in Pabpn1l +/+ and Pabpn1l -/- oocytes and embryos. (A) Poly(A) tail assay results of indicated transcripts documenting the length of poly(A) tail in Pabpn1l +/+ and Pabpn1l -/- oocytes and embryos. The poly(A) tails of Gapdh are used as an internal control. Each sample was prepared from 100 oocytes or embryos. (B) Length distribution of poly(A) tails. The Y-axis represents averaged relative signal intensity, and the X-axis represents the length of the PCR products based on their electrophoretic mobility.

## Discussion

PABPs are highly conserved eukaryotic proteins, which specifically recognize the poly(A) sequence of mRNA 3’-end. They are widely involved in biological processes and display clear functional differences. These differences may be due to distinct cellular and subcellular localization of PABPs and variations in the structure of these proteins and interacting molecules. PABPN1L belongs to the class of PABPs distributed in the nucleus and having only one RRM domain. Early in the 21st century, its Xenopus homologous protein ePABP2 was considered to have the expression pattern of a maternal factor. Since then, it has been found that ePABP2 can bind AAAA nucleic acid sequence, similarly to PABPN1^31^. However, the function of PABPN1L in female gametes of Xenopus and mammals has not been identified yet.

Pabpn1l -/- mice, a model of abnormal splicing of Pabpn1l, and a gene trapping model, Pabpn1l tm1a/tm1a, were used in this study. It was found that PABPN1L has typical characteristics of a maternal factor, and its deletion leads to early embryo block without affecting oocyte maturation and fertilization. Similar phenotypes were reported in a contemporaneous study ^32^. We found that thousands of transcripts were altered in MII oocytes in the absence of Pabpn1l, an effect similar to that of the Btg4 knockout. Because of the abnormal deadenylation of mRNA in Btg4-/- MII oocytes^12^, the poly(A) length of maternal mRNA in Pabpn1l -/- oocytes was assessed. Timm9, Eloc, Elobl, and Fbxw13 mRNAs in the fertilized Pabpn1l -/- eggs showed shortened poly(A) tail. Surprisingly, the poly(A) tail of these mRNAs was significantly shorter in the MII oocytes of Pabpn1l -/- than of Pabpn1l +/+. In another study, although the authors of that report did not describe the results of the PAT assay in the MI phase oocytes, it was evident that the poly(A) tail of Zp2, Nobox, Oosp2 and Ube2t mRNAs in the MI phase oocytes with Pabpn1l deficiency was significantly shorter than in the control group^12^. This phenomenon of poly(A) tail shortening at the MII stage was present not only in the upregulated mRNA of Pabpn1l -/- MII oocytes but also in Cdc123 mRNA and Gapdh mRNA, with no change in the total amount. Conversely, the mRNA of Zp2, Nobox, Oosp2, and Ube2t in Btg4 -/- MII oocytes had longer poly(A) tail. Interestingly, despite significant shortening of the poly(A) tail, oocytes could not start the degradation process at the MII stage in Pabpn1l -/-. These results suggest that PABPN1L may play a role in antagonizing poly(A) tail shortening in oocytes, independently of its involvement in the process of maternal mRNA degradation. How this maternal mRNA with shortened poly(A) escapes degradation in MII oocytes is be a question that can be explored in the future. In another study, Vieux and collaborators documented that lengthening the poly(A) tail could not inhibit mRNA degradation during oocyte maturation^33^. This finding supports the possibility that the poly(A) tail length of some mRNAs does not affect its degradation during oocyte maturation. One possible hypothesis is that PABPN1L has a function similar to that of PABPN1, which can promote the growth of poly(A) tail of mRNA; alternatively, PABPN1L is necessary to recruit molecules for mRNA degradation. Therefore, after the deletion of Pabpn1l, the poly(A) of maternal mRNA is shortened, but the degradation process is terminated.

The degradation of maternal mRNA affects female reproductive outcomes. Although IVF and ICSI have helped a lot of infertile couples, early embryo block can occur, leading to treatment failure^34–36^. Several genes are involved in oocyte maturation arrest (such as TUBB8^37^ and PATL2^38^) and fertilization failure (such as TLE6^39^ and WEE2^40^). Recent studies demonstrated that BTG4 variants could be used as a genetic marker for zygotic cleavage failure (ZCF)^41^. Our study provides new clues to understanding the mechanism of ZCF. By analyzing the gnomAD data, we found that there are low-frequency PABPN1L variants in the population, and these variants can result in abnormal splicing of PABPN1L. Moreover, by simulating the splicing abnormality of intron1, we found that in the Pabpn1l -/- female infertility, oocytes were blocked in the 1-cell to 2-cell stage after fertilization. Our study provides a new genetic diagnostic target for human ZCF.

## Materials and Methods

### Animals

Pabpn1l -/- mice were generated as described below. Pabpn1l tm1a(EUCOMM)Wtsi/+ (referred to as Pabpn1l tm1a/+) mice were obtained from the CAM-SU Genomic Resource Center of Soochow University. The null allele was generated using JM8A3.N1 ES cells (Genome Research Limited, Wellcome Trust Sanger Institute). The L1L2_Bact_P cassette was inserted at position 122622344 of Chromosome 8 upstream of Pabpn1l exon 2. The mice were maintained and used in experiments according to the guidelines of the Institutional Animal Care and Use Committee of Nanjing Medical University (China). The animal use protocol has been reviewed and approved by the Animal Ethical and Welfare Committee (Approval No. IACUC-1912001).

### Quantitative RT-PCR assay

The samples include somatic tissues, oocytes, and preimplantation embryos from wild-type, Pabpn1l-/- and Pabpn1l tm1a/tm1a. Total RNA was extracted from the samples using TRIzol reagent (Thermo Fisher Scientific, Carlsbad, CA, USA). The concentration and purity of RNA were determined using a NanoDrop 2000C (Thermo, Waltham, MA, USA)absorbance at 260/280 nm. Total RNA (1 μg) was reverse transcribed using a HiScript II Q RT SuperMix (Vazyme, R222, Nanjing, China) according to the manufacturer’s instructions. The cDNA(dilution 1:4) was then analyzed by quantitative RT-PCR in a typical reaction of 20 μl containing 250 nmol/l of forward and reverse primers, 1 μl cDNA and AceQ qPCR SYBR Green Master Mix (Vazyme, R222, Nanjing, China). The reaction was initiated by preheating at 50°C for 2 min, followed by 95°C for 5 min and 40 amplification cycles of 10 s denaturation at 95°C and 30 s annealing and extension at 60°C. Gene expression was normalized to 18 s within the log phase of the amplification curve. The primer sequences are listed in supplementary Table S2.

### Pabpn1l-EGFP mRNA *in Vitro* transcription and microinjection

The mouse Pabpn1l coding sequence (CDS) corresponding to full-length Pabpn1l was PCR-amplifified from a mouse ovarian cDNA pool, and EGFP tag sequences were inserted before stop codon using Phanta Max Super Fidelity DNA Polymerase (Vazyme, P505,Nanjing, China). The PCR products were purifified using the FastPure Gel DNA Extraction Mini Kit (Vazyme, DC301-01, Nanjing, China) and then digested by BamHI and HindIII restriction enzymes (New England Biolabs, Inc.). C-terminal EGFP-tagged mouse full-length Pabpn1l was cloned into pcDNA3.1(+) vectors by the ClonExpress MultiS One Step Cloning Kit (Vazyme, C113). To prepare mRNAs for microinjection, expression vectors were linearized and in vitro transcribed using the mMESSAGE mMACHINE T7 ULTRA Transcription Kit (Thermo Fisher Scientific, Carlsbad, CA, USA). Images were captured using an LSM800 confocal microscope (Carl Zeiss AG, Jena, Germany).

### Genotyping

Tail clips were subjected to standard DNA-extraction procedures. The Pabpn1l-/- gene was identified via PCR using the Pabpn1l-For and Pabpn1l-Rev. For Pabpn1ltm1a/+, the forward primer was Pabpn1l-F, the reverse primer was Pabpn1l-KO-R and Pabpn1l-WT-R. The products were used in a PCR reaction with genespecific primers under the following conditions: 30 s at 95 °C, 30 s at 58 °C, and 30 s at 72 °C. PCR products were analyzed on a 2.0% agarose gel. The primer sequences are listed in supplementary Table S2.

### fGeneration of Pabpn1l-/- mice by CRISPR/Cas9

Cas9 mRNA and single guide RNAs (sgRNAs) were produced and purified as previously described^43–47^. In brief, the Cas9 plasmid (Addgene, Watertown, MA, USA) was linearized by restriction enzyme digestion with AgeI and then purified using a MinElute PCR Purification Kit (Qiagen, Duesseldorf, Germany). Cas9 mRNA was produced by in vitro transcription using a mMESSAGE mMACHINE T7 Ultra Kit (Ambion, Austin, TX, USA) and purified using a RNeasy Mini Kit (Qiagen, Duesseldorf, Germany) according to the manufacturer’s instructions. The sgRNAs were designed on the basis of intron1-exon2 of Pabpn1l. The target sequence of sgRNA was 5’-GCCACCCTGTGTACAGAGAAAGG-3’ and 5’ - GGCCCACAGTTTCAGCTTGATGG-3’, respectively. The sgRNA plasmid was linearized with DraI and then purified using a MinElute PCR Purification Kit (Qiagen, Duesseldorf, Germany). sgRNA was produced using the MEGA shortscript Kit (Ambion, Austin, TX, USA) and purified using the MEGA clear Kit (Ambion, Austin, TX, USA) according to the manufacturer’s instructions. Cas9 mRNA and sgRNA were injected into mouse zygotes obtained by mating of wild-type C57BL/6 males with C57BL/6 superovulated females.

### Fertility test

Adults each genotype were subjected to fertility tests. Each male was mated with three wild-type mice, and the vaginal plug was checked every morning. The dates of birth and number of pups in each litter were recorded.

### Histological analysis

Ovaries were fixed in 4% PFA overnight at 4 °C, and then dehydrated through a series of graded ethanol solutions and xylene before paraffifin embedding. Serial sections (5 μm) were attached to slides, heated at 60 °C, followed by washing in xylene and rehydration through a graded series of ethanol and double-distilled water and then staining with Hematoxylin and Eosin (HE) reagents.

### Superovulation and IVF

For superovulation, female mice (21–23 days old) were intraperitoneally injected with 5 IU of pregnant mare’s serum gonadotropin (PMSG) (Ningbo Sansheng Pharmaceutical Corporation, Zhejiang, China). After 44 h, mice were then injected with 5 IU of human chorionic gonadotropin (hCG) (Ningbo Sansheng Pharmaceutical Corporation, Zhejiang, China). Eggs collected from superovulated females 14 h after hCG injection were placed in 150 μL of HTF medium, drop covered with paraffin oil. Epididymal sperm were collected from 3-mo-old male mice and incubated in HTF (AIBI Bio-Technology Co.Nanjing, China) medium for 40min for capacitation. Capacitated sperm were added to the drop containing eggs at a final concentration of 2 × 10^5^ sperm/ml. After 4 h of coincubation, the formation of pronuclei was observed. The fertilized eggs were blown clean and cultured in KSOM (AIBI Bio-Technology Co.Nanjing, China) medium. Images were captured using an Ti2-U microscope (Nikon, Tokyo, Japan).

### RNA-seq and data analysis

GV oocytes, MII oocytes and zygotes were collected and mixed from three mice of each genotypes seperately. Each group contained a total of 30 oocytes or zygotes. Each cell sample after sorting can be stored in 6μl SMART-Seq™ v4 kit lysate. The volume of fresh cells was measured, and SMART-Seq™ v4 Ultra™ Low Input RNA Kit for Sequencing (Clontech) was used for cell lysis and first strand cDNA synthesis. Then, the first stand cDNA was amplified by LD-PCR. The amplified double stranded cDNA was purified by AMPure XP beads. The PCR products were purified by AMPure XP beads and the final library was obtained. Using Agilent 2100 Bioanalyzer to detect the insert size of the library. After the library was qualified, the library was pooled according to the effective concentration and the demand of target offline data, and was sequenced by Illumina platform. Then quality control was done using FASTQC. Raw reads were trimmed with Trimmomatic (v0.39) and mapping to mouse genome (GRCm38),using Hisat2(v2.2.0) (Daehwan et al., 2015). Only the uniquely mapped reads were kept for the subsequent analysis. This was followed up with Stringtie for transcript assembly. the numbers of reads per gene were normalized via performing TMM (trimmed mean of M values) normalization in the Bioconductor package edgeR and different genes were accessed at a P value of <0.05, false discovery rate (FDR) of <0.05 and fold change (FC)| of >2. The Spearman correlation coefficient (rs) was calculated using the cor function in R. Gene Ontology (GO) enrichment analysis were performed by ClusterProfiler package in R.

### Poly(A) tail (PAT) assay

Total RNA was isolated from oocytes at the indicated stages with an RNeasy Mini kit (Qiagen, Duesseldorf, Germany). The Poly(A) Tail-Length Assay Kit (Affymetrix, Cleveland, Ohio,USA) is designed for G/I tailing up to the total RNA. All necessary components are provided to perform 4 reverse transcription and 16 PCR reactions on each tail-extended samples. Reaction products are then assessed by gel electrophoresis. Bands were quantified using the “Plot profiles” function of ImageJ software, normalized using the maximum signal intensity in each lane, and the averaged values of three biological replicates were plotted.

### Statistical analysis

All experiments were repeated at least three times. The differences between treatment and control groups were analyzed using one-way ANOVA or unpaired two-tailed t-tests. P-values < 0.05 were considered to indicate statistical significance. All data represent the mean ± the standard deviation of the mean. Analyses were performed using the Microsoft Excel or GraphPad Prism 6.0.

## Acknowledgements

We thank Lu Liu, Qianneng Lu, Xiaoning Yu, Yue Hu, Shuai Zhou, Mingrui Li and Ran Huo for their help on ART, Yiqiang Cui for the sgRNA design. This work was supported by the National Key Research and Development Program of China 2016YFA0500902 (to M.L.); Natural Science Foundation of China (81701446 to Y.W. and 31771654 to M.L.); the Natural Science Foundation of Jiangsu Province (Grants No. BK20190081 to M.L.); and Qing Lan Project (to M.L.).

## Authors’ roles

Y.W., T.F., and X.S. performed most of the experiments; S.L., Z.W., X.Z., J.Z. and S. Z. performed some of the experiments; Y.W., T.F., S.L., Z.W. and M.L. analyzed the data; Y.W. and M.L. initiated the project and designed the experiments; Y.W., T. F., X.L., and M.L. wrote the paper. All authors have read and approved the final manuscript.

